# Association of a lincRNA *postmortem* with suicide by violent means and *in vivo* with aggressive phenotypes

**DOI:** 10.1101/257188

**Authors:** Giovanna Punzi, Gianluca Ursini, Giovanna Viscanti, Eugenia Radulescu, Joo Heon Shin, Tiziana Quarto, Roberto Catanesi, Giuseppe Blasi, Andrew E. Jaffe, Amy Deep-Soboslay, Thomas M. Hyde, Joel E. Kleinman, Alessandro Bertolino, Daniel R. Weinberger

## Abstract

**Objective:** Previous findings suggest that differences in brain expression of a human-specific long intergenic non-coding RNA (*LINC01268*; GRCh37/hg19: *LOC285758*) may be linked to aggressive behavior and suicide. The authors sought to replicate and extend these findings in a new sample, and translate the results to the behavioral level in living healthy subjects.

**Method:** The authors examined RNA sequencing data in human brain to confirm the prior postmortem association of the lincRNA specifically with suicide by violent means. In addition, they used a genetic variant associated with *LINC01268* expression to detect association with *in vivo* prefrontal physiology related to behavioral control. They finally performed weighted gene co-expression network analysis (WGCNA) and gene-ontology analysis to identify biological processes associated with a *LINC01268* co-expression network.

**Results:** In the replication sample, prefrontal expression of *LINC01268* was again higher in suicides by violent means (N=65) than both non-suicides (N=78; 1.29e-06) and suicides by non-violent means (N=46; p=1.4e-06). In a living cohort, carriers of the minor allele of a SNP associated with increased *LINC01268* expression in brain scored higher on a lifetime aggression questionnaire and show diminished engagement of prefrontal cortex (BA10) when viewing angry faces during fMRI. WGCNA highlighted the immune response.

**Conclusions:** These results suggest that *LINC01268* influences emotional regulation, aggressive behavior and suicide by violent means; the underlying biological dynamics may include modulation of genes potentially engaged in the immune response.

---

Suicide is the 10th leading cause of death in the United States for all age groups combined(1), a trend steadily increasing in the last 15 years against the backdrop of generally declining mortality(2). Psychiatric disorders, while strongly predictive of suicidal ideation, are less suitable in predicting who will actually act on such thoughts; in contrast, severe anxiety/agitation and poor impulse control are better indicators of higher risk for suicide plans and attempts(3). The choice of violence as a means of suicide has been considered a correlate of the cumulative amount of lifetime impulsive-aggressive behavior(4) and, consistently, an association between violent offending and suicide has been suggested as well(5). The two phenomena can ultimately coexist in a single episode of acting-out (homicide-suicide or ‘extended suicide’)(6). Recent findings(7) also show that the choice of a violent method for a self-harm episode translates into a remarkably higher risk for suicide in the short term; current research suggests an increase in such self-inflicted injuries particularly among youth(8).

As suicidal ideation is a poor predictor of outcome, the detection of biological markers for violent suicidal risk represents an important insight for the development of prevention approaches(9) as well as for understanding aggressive behavior itself. Genetic factors for traits that render an individual prone to engage in drastic measures such as suicide have been broadly studied(10), with impulsive aggression as a strong candidate intermediate phenotype(11, 12). One approach in this line of research has been to investigate genetic signaling involved in the action of drugs credited as effective against suicide, such as lithium(13). Intriguingly, levels of lithium in drinking water have been associated with lower suicidality(14, 15) and even with lesser rates of homicides(16). In a prior study of a small cohort of schizophrenic patients(17), we found that the expression of *MARCKS*, a gene implicated in suicidal behavior(9, 18, 19) and down-regulated by lithium(20), is significantly increased in the dorsolateral prefrontal cortex (DLPFC) of suicides specifically by violent means (i.e. a suicide where self-harm with lethal intent is carried out by actively inflicting painful injuries; details in Method). *MARCKS*’ association to suicide was in fact statistically conditioned on a lincRNA (long inter-genic non-coding RNA) that maps adjacent to it: *LINC01268* (GRCh37/hg19: *LOC285758*)(17).

Intriguingly, the non-protein-coding portion of the human genome has been increasingly recognized as a source of evolutionary organismal complexity(21). In this regard, evolutionarily conservation annotations of lncRNAs (e.g., phyloNONCODE) show that *LINC01268* is poorly conserved in rodents, but highly conserved in non-human primates and, notably, transcribed only in humans. While molecular mechanisms lie in the chain of causation from genome to behavior, aggressive and suicidal acts ultimately involve complex, human-specific behavior and emotional control, and indeed long-term risk for suicide at the level of higher-order brain function has been related to control dysfunction of prefrontal cortex(22).

The main hypothesis underlying this report is that the choice of a violent method of committing suicide represents a more precise feature to specifically target in order to detect genetic signatures for this extreme behavior in general. The aims of this study are: 1) to replicate our earlier finding of a link between expression of *LINC01268* in prefrontal cortex and suicide specifically by violent means(17) by analyzing a new sample of human brains, including patients with schizophrenia, depression and bipolar disorder; 2) to translate this finding into a behavioral context via a SNP associated with *LINC01268* expression and a fMRI paradigm targeting prefrontal function related to negative emotion processing; and 3) to further explore the biological meaning of the transcript through analysis of the network of genes co-expressed with it using WGCNA (Weighted Gene Co-Expression Network Analysis).

## METHOD

### Postmortem Data

#### Subjects

A large cohort of brains with a RIN ≥6.9 was obtained from the Lieber brain repository, focusing on the available RNA-sequencing dataset from DLPFC (BA46/9)(23). The relationship between suicide and the lincRNA was first explored in a diagnostically mixed sample of 189 adult Caucasian patients (PZ) who met DSM-IV criteria for a lifetime diagnosis of schizophrenia (SCZ), bipolar disorder (BPD) or major depressive disorder (MDD), individuals not included in our earlier study. The comparison between suicides and non-suicides is best evaluated in patients only, to control for associations with a diagnostic state. A single race (Caucasian) was chosen to avoid confounding effects on gene expression based on racial genomic variation; only donors ≥13 years were selected, since gene expression displays strong non-linear changes during developmental stages(24). One brain donor was discarded as an extreme outlier for *LINC01268* expression. Within this replication sample, 111 individuals completed suicide and 78 were non-suicidal. Within the suicide cohort, 46 were suicides by non-violent means, 65 suicides by violent means. In addition to testing for association between suicide by violent means and our lincRNA in this new sample, we also analyzed the results after adding patients from the previous study. Following the described criteria for inclusion, the total brain sample comprised 228 subjects. Table S1 provides a summary of the combined patient sample. ST1a shows that suicides are younger than non-suicides (p=0.002) especially suicides by violent means (p<0.002), and that among suicides, more men choose violent means than females (p<0.006). These data support the involvement of both age-and gender-related factors in impulsive/aggressive behaviors.

#### Postmortem Brain Tissue

Postmortem brains from the LIBD were collected at autopsy from several medical examiners; some of these brain tissues were collected at the NIMH. A similar uniform procedure was employed in the acquisition, processing and dissection of all samples regardless of source and affiliation; detailed methods relating to the collection have been reported elsewhere(23, 25).

#### Manner of Death and Suicide

Cause and manner of death and contributory causes or medical conditions related to death were obtained from medical examiner documents(23). Non-suicidal deaths included natural deaths, accidents and homicides. In regard to the suicide group, we expect that people who were generally aggressive and prone to physical violence would, in the course of taking their own lives, tend to choose a violent method. We consider ‘suicide by violent means’ a suicide where self-harm with lethal intent is carried by actively inflicting painful injuries. Examples are: hanging, gunshot, blunt force, sharp force, jumping from heights, motor vehicle accident et cetera. On the other hand, a ‘suicide by non-violent means’ consist of actions that are ‘physiological’, like swallowing or drinking something that is not hurting at that moment (i.e. drug overdose) do not require the subject to produce injuries on his/her body to die. The same applies to breathing in a *suicide bag* (see below), or in a confined space full of a toxic gas like carbon monoxide, both known to make exposed individuals unconscious with minimal discomfort. Such methods represent the first choice of those who do not want to suffer, and, instead, desire to black out as soon as possible and never wake up. The choice of a specific method likely reflects the interplay of multiple determinants, including the availability of a particular suicidal means; however, at the individual level, preferences toward one or the other group appear to influence the ultimate pattern of choices. It has been noted(26) that when access to a suicide method is restricted and if, as a consequence, a substitution occurs, the transition probability between two similar methods (i.e. from firearm to hanging, both violent) is much higher than between for example poisoning and a violent method. Several studies have attested to such peculiar phenomenon(27–32).

For accuracy, we do not simply rely on the medical examiner labelling (e.g. ‘poisoning’, or ‘asphyxia’), but on a case-by-case in depth evaluation of all available information, including interviews with next of kin. Suicidal poisoning by ingesting, for example, caustic soda, which gives burns and extreme pain since the first sip, is actually violent. Similarly, in some cases where the final cause of death is indeed poisoning (i.e. pills), other actions can also be taken by the victim to inflict serious injuries, eventually making the method violent. Additionally, many suicides classified under the generic term of ‘asphyxia’ may fall in either category. The circumstances under which deaths by asphyxia occur span both more active, violent interventions (such as smothering by stuffing one‘s airways with foreign objects), and peaceful settings like a *suicide bag*: a DIY device (a.k.a ‘the exit bag’ among suicide supporters), consisting of a drawstring plastic bag inflated with an inert gas like helium, which prevents alarm responses, leading to a painless death. Another suicidal manner that deserved consideration, not being intuitively classifiable, is drowning. In contrast with previous literature(33), we do not categorize drowning *a priori* as a nonviolent method. Suicides by drowning may, in fact, occur by jumping from heights into the water and be ultimately caused by the blunt-force impact. In our opinion, this approach reflects more a violent drive as opposed to the predisposition underlying a drowning occurring, for instance, in a bathtub while severely intoxicated. We excluded cases where manner of death was pending or not determined at the time of the curation, and suicidal samples with ambiguous, or indefinable means of suicide in regard to the level of violence employed. Among the remaining suicides, most deaths distinctly fell within the violent or non-violent category. When this was not obvious, an in-depth assessment was obtained using detailed narrative summaries based on all available sources of historical information(23). Table 1 provides details about the suicides in the whole sample of 228 subjects (see also ‘Subjects’). Attribution of suicidal method was determined blind to the postmortem data.

**Table 1:** Manners of suicide.

| ***Suicides by Violent Means*** |  | ***Suicides by Non-Violent Means*** |  |
| --- | --- | --- | --- |
| Hanging | 38 | Drug poisoning | 31 |
| Gunshot | 15 | Overdose | 13 |
| Blunt force | 13 | CO poisoning | 4 |
| Sharp force | 5 | Asphyxia<sup>2</sup> | 1 |
| Asphyxia<sup>1</sup> | 4 | Asphyxia<sup>3</sup> | 1 |
| Motor vehicle accident | 1 |  |  |
| Low voltage electrocution | 1 |  |  |

<sup>1</sup>Violent method; <sup>2</sup>Suicide bag; <sup>3</sup>Positional (due to drug intoxication)

#### Gene Expression

RNA Extraction, sequencing and data processing were performed as previously described(34) details in SI.

#### Genotyping

Genomic DNA was extracted from cerebellum of the same samples using standard procedures with Flexi Gene DNA kits (Qiagen, Germany). Genotyping was performed as previously described(23, 35, 36), and an independent set of SNPs obtained by LD-pruning(37) was used to perform genome-wide clustering to obtain multidimensional scaling (MDS) components for quantitative measures of ancestry. For our eQTL analysis, we removed SNPs showing minor allele frequency <15%, genotype missing rate >5%, or deviation from Hardy-Weinberg equilibrium (p<0.0001). Additional QC was performed on individual genotyping results. Individuals were removed if their overall genotyping rate was below 97%. The data were checked for sample duplications and cryptic relatedness.

#### eQTL Identification

We searched for single nucleotide polymorphisms (SNP) associated with *LINC01268* expression in a sample of healthy subjects (HSs, N=105) from the Lieber repository, in order to avoid confounding effects of diagnosis and treatment. Similarly to the PZ sample, these subjects were both Caucasian and ≥13 year old, for design homogeneity. The age criterion was further supported by the evidence that the brain expression of the lincRNA did not actually show non-linear changes above the age of 13 years in the HSs cohort (SF1).

#### WGCNA

Weighted Gene Co-Expression Network Analysis(38) was performed on RNA-seq data from suicide by violent means, with Gene Ontology (GO) analysis on each module identified. Validation of the modules was further achieved with module preservation. This method is further detailed in SI.

### Behavioral Data

#### Subjects

72 Italian healthy adults (all >18 years), unrelated, Caucasians, non-offenders, genotyped for rs774796, were studied. One outlier for adulthood aggressiveness (‘Adult BG’, see below) was removed and two more subjects were excluded since the aggressive behaviors uncovered by the interview did not fully qualify them as non-offenders. The final sample consisted of 69 subjects (males 32; age=26.12, SD±4.84) of whom 51 were TT, 13 CT and 5 CC for the genotype of interest. 58 individuals from this cohort underwent 3 Tesla fMRI. They were matched for age, gender, handedness (Edinburgh Inventory), socio-economical status (Hollingshead Scale) and IQ (WAIS-R).

#### Recruitment

Healthy volunteers recruited by the Group of Psychiatric Neuroscience at the University of Bari, Italy, entered the behavioral study. Participants provided informed consent in writing; protocols and procedures were approved by the local Institutional Review Board. All subjects underwent the Structured Clinical Interview for *DSM-IV* to exclude Axis I psychiatric disorders. None of them had a history of significant drug or alcohol abuse (no active drug use in the 6 months prior to the study), head trauma with loss of consciousness, or significant medical illness.

#### DNA Extraction and Genotyping

DNA was extracted from whole blood using standard procedures, with QIAamp DNA Blood Midi Kit (Qiagen, Germany), and genotyped as described for the postmortem sample.

#### Life History of Aggression

All the subjects were evaluated by a psychiatrist using the Brown-Goodwin (BG) Questionnaire, in its revised version consisting of 11 items(39), which measures actual behavioral manifestations rather than temperament. The interview provides distinct scores for each phase of life (≤12 years, 12-18 years and ≥18 years, i.e. adulthood) and a total score. For the purposes of this study, only the score related to adulthood (‘Adult BG’; mean value 15.38, SD±3.16) was analyzed, as an index of current aggressiveness.

#### fMRI paradigm

The event-related fMRI paradigm(40) consisted of presentation of faces with angry, fearful, happy and neutral emotional expressions from a validated set of facial pictures (NimStim, www.macbrain.org/resources.htm)(41). Subjects were asked whether they would ‘approach’ or ‘avoid’ the faces presented during the task (explicit emotional processing(40)). In the present study, we focused on brain responses during processing of angry faces (angry stimuli *vs* crosshairs), since they are considered signals of threat and tend to evoke hostile feelings(42). *fMRI data acquisition*. Details are reported in SI.

### Statistical Analysis

All data processing was performed using the ‘R’ statistical language(43). Grubbs tests and 3xInterquartile Range method(44) were used to identify outliers.

Normality Test (Shapiro-Wilk) showed that *LINC01268* expression did not exhibit a normal distribution; therefore, logarithmic values (log2) were calculated and employed in each analysis, with an offset of 1 to avoid issues with 0s. Two sample t-test and *X*_2_ test for differences were used to compare demographic data across manners of death in the postmortem cohort and across genotypes in the Italian sample. *X*_2_ test was also used to test the effect of comorbidity on suicide by violent means *vs* non-suicide. Linear Models were used for comparisons of mRNA expression between groups, with manner of death as an independent variable (or comorbity in the sensitivity analyses) and with age, sex and RIN as covariates (age and RIN in the analysis by sex). Linear Models were used for the eQTL analysis, with 10 expression heterogeneity PCs and 3 genetic PCs as covariates(36). Linear Models were used to investigate the relationship between Adult BG and genotype, covarying for age and sex. *fMRI data analysis, WGCNA*: details are reported in SI.

## RESULTS

### Postmortem prefrontal expression of *LINC01268* and suicide

To evaluate the relationship between the DLPFC expression of the lincRNA and suicidal behavior in general in a new sample of subjects (N=189), two main groups of suicidal and non-suicidal brain donors were first compared. The analysis (SF2a) confirmed the association of *LINC01268* mRNA expression levels (RPKM) with suicide, in that people who took their lives had significantly higher levels of *LINC01268* in their DLPFC than non-suicidal donors (t=2.750, p=0.006). Subsequently, the subgroup of suicides by violent means was compared to non-suicides and suicide by non-violent means. This analysis (SF2b) confirmed that the association between *LINC01268* and suicide was driven exclusively by the suicides by violent means: the violent suicide group had higher *LINC01268* expression in DLPFC than both non-suicides and suicides by non-violent means (t=5.064, p=1.29e-06; t=5.116, p=1.4e-06, respectively). Non-suicides and suicides by non-violent means did not differ in *LINC01268* expression values (t=-0.505, p=0.614). The analysis yields similar results by adding ‘diagnosis’ to the model as a covariate. Figure 1a, b displays our cumulative findings by including also the subjects that we have previously studied(17): the combined sample comprises brains from 228 individuals, thus increasing statistical power.

**Figure 1:**
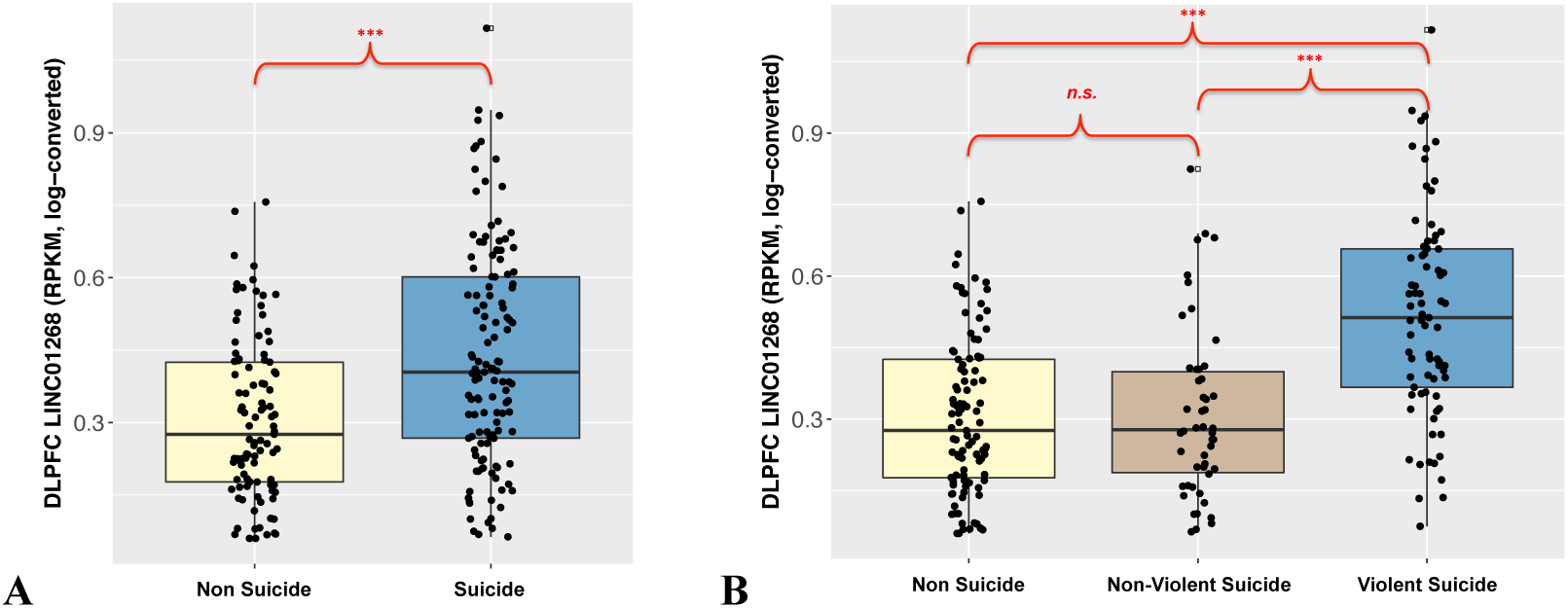
Boxplots of the effect of manner of death on the DLPFC expression of *LINCO1268* in the total sample (N=228, All PZ, with age, sex and RIN as covariates). Manner of death is significantly associated with *LINC01268* expression, which is greater in suicide completers compared with non-suicidal deceased **(A)**, and specifically in suicides by violent means (N=77) compared with non-suicides (N=101) and suicides by non-violent means (N=50) **(B)**. Non-suicides and suicides by non-violent means do not differ. Statistics: **(A)** non suicide vs suicide: t=3.979, p=9.37e-05 (df 1, 223); **(B)** non suicide vs violent suicide: t=6.527, p=7.16e-10 (df 1, 173); non suicide vs non-violent suicide: t=0.056, p=0.955322 (df 1, 146); nonviolent suicide vs violent suicide: t=5.450 p=2.66e-07 (df 1, 122).

We further tested this larger sample in a number of sensitivity analyses. Additional boxplots show the effect of suicide by violent means *vs* non-suicide on the DLPFC expression of *LINC01268*, divided by sex and diagnosis (SF3 and SF4). The expression of the lincRNA is significantly higher in suicide by violent means than in non-suicide in each sex (SF3), and in each diagnosis (SF4), although in the BPD group there is only a statistical trend, likely due to the relatively smaller size of this sub-cohort (see ST1). Moreover, a co-diagnosis of alcohol and/or substance use is not associated with suicides by violent means (*X*_2_=1.0789, p=0.299); nor is it associated with *LINC01268*’s brain expression (t=-0.917, p= 0.3604). Furthermore, to minimize artifacts such as occult RNA quality differences that may affect the results and partially explain the differences between the three groups, we repeated the analysis with the addition of adjusting covariates from a PC analysis, as reported in SI (*‘Additional Sensitivity Analyses’*). These analyses, confirming all the results, excluded also the potential effect on gene-expression of the injuries resulting from the violent action.

### Rs7747961 as a candidate eQTL for *LINC01268*

We searched for cis-acting eQTLs possibly associated with expression of the *LINC01268* transcript, by evaluating SNPs within 100,000 bp upstream and downstream of the gene. Of the SNPs tested in a dataset from brains of HSs, none survived correction for multiple comparisons across the genome (FDR); however, SNPs with nominally significant p-values (<0.05) were further explored. An additive model uncovered five SNPs that fit this criterion: three of them are independent in our sample (r2<0.1, rs503593, rs9481396 and rs7747961); two others were in strong LD with rs9481396 (rs73547709 [R^2^=0.964, D’=1.000] and rs59034807 [R^2^=0.892, D’=0.962]). While rs9481396 showed overdominance, thus not allowing us to identify the allele clearly associated with increased expression, the associations of rs503593 and rs7747961 with gene-expression had more linear distributions (p<0.05). In further analysis employing a dominant model, only rs7747961 showed nominally significant association with *LINC01268* levels (p=0.041): carriers of the minor C allele (CT and CC genotypes) showed significantly higher values of *LINC01268* transcript than homozygotes for the major allele (Figure 2a). Rs7747961 lies 2205 bases upstream of the MARCKS TSS and is linked with regulatory element markers at the SNP site in ChIP experiments from publicly available datasets from brain (e.g. www.broadinstitute.org/mammals/haploreg/haploreg.php). As this SNP did not reach multiple testing-corrected levels of significance, we performed additional sensitivity analyses. We selected the 23 linkage disequilibrium (LD)-independent SNPs within the 200kb window of our original SNP expression analyses using pruning of the genotype data (LD R^2^<0.1)(37). Rs7747961 had stronger eQTL statistics than all 23 independent SNPs (p-value range: 0.1-0.9) in the discovery dataset. Lastly, we tested rs7747961 as a potential eQTL for *LINC01268* in DLPFC in the independent CommonMind Consortium (CMC)(45) control sample dataset (N=216), and confirmed this association, which was also directionally concordant (in both additive and dominant models, p<0.05). Hence, rs7747961 was considered suitable as an eQTL for interrogation of a potential proxy of *LINC01268* expression in *in vivo* clinical genetic studies.

**Figure 2:**
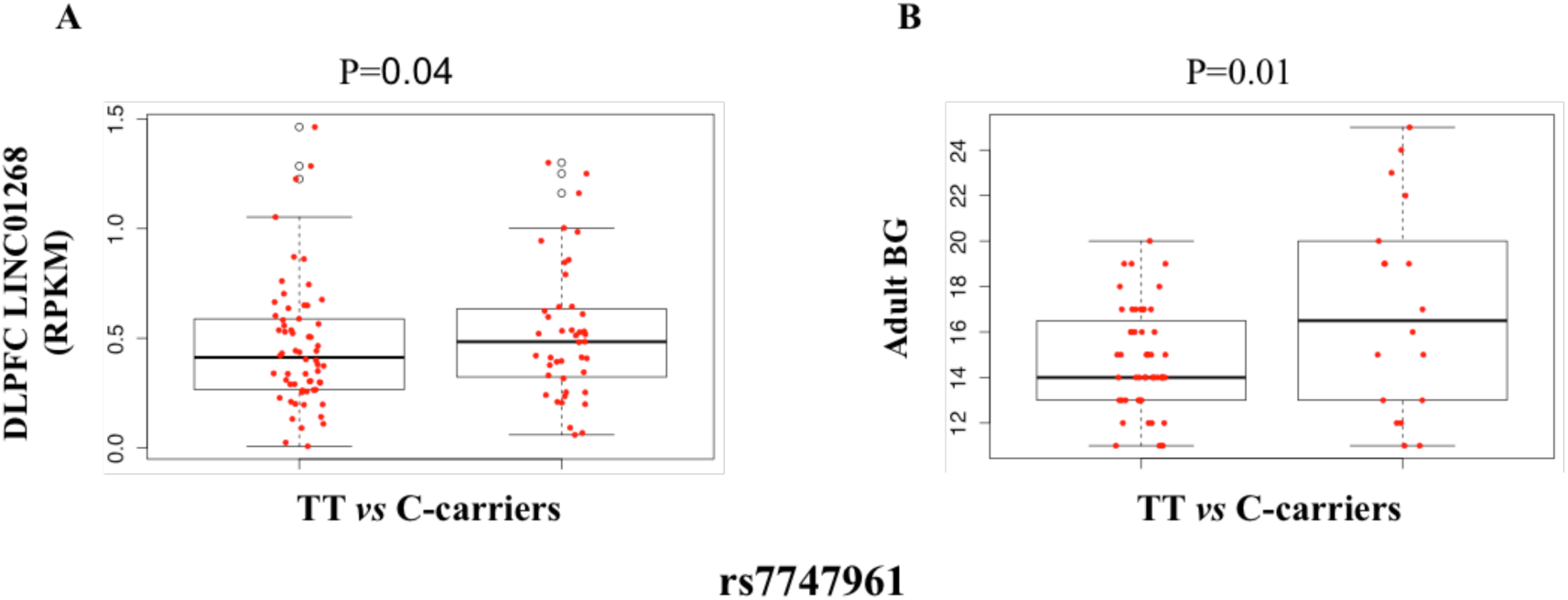
**(A)** Boxplots of the association of rs7747961 genotype with DLPFC expression of *LINC01268*. Statistics for eQTL LM (10 expression heterogeneity PCs and 3 genetic PCs) are reported at the top of the graphic. Both in additive (not shown) and dominant models, rs7747961 genotype is nominally significantly associated with *LINC01268* expression, which is greater in C-carriers (right) compared with homozygotes for the major allele T (left). **(B)** Boxplots of the effect of rs7747961 genotype on levels of adult ratings of aggressiveness (Adult BG). Statistics for LM (with age and sex as covariates) are reported at the top of the graphic. Rs7747961 genotype is significantly associated with adulthood aggressiveness, which is greater in C-carriers (right) compared with homozygotes for the major allele T (left).

### Association of rs7747961 with behavioral measures of aggressiveness

To translate *LINC01268* transcription to the behavioral level, and expand our inquiry to living subjects, an independent sample of living HSs was assessed for a trait measure of aggressiveness and genotyped for rs7747961. The relationship between Adult BG and genotype was tested using the same dominant model used with the postmortem expression results. Remarkably, the same genotypes (the carriers of the C allele) associated with increased expression of *LINC01268* in postmortem brains were also associated with more aggressiveness in healthy subjects (p=0.01, Figure 2b), with the same allelic directionality. We further considered the 23 LD SNPs described above, and found that rs7747961 was more strongly associated with Adult BG than any of them (p-value range: 0.1-0.9), suggesting that this SNP may play a role in brain function. Finally, in the largest sample exploring children’s aggressive behavior to date (N=18,988) and the first GWAS on the subject, the minor allele for rs7747961 was consistently associated with higher aggression in children, although with a p-value below the threshold for GWA significance (FDR p_one-sided_<0.05)(46).

### Rs7747961, aggressiveness and brain activity during explicit emotion processing

Given the association between *LINC01268* mRNA brain levels and rs7747961, and between the latter and aggressiveness, data from an fMRI study of response to emotionally charged stimuli (i.e., angry faces) in the HS sample were analyzed. The factorial regression analysis indicated no main effect of rs7747961 or Adult BG on the imaging data, but uncovered an interaction between rs7747961 and Adult BG in right BA10 (MNI coordinates: x=36, y=50, z=28; Z=3.53; P_FWE-corr_.=0.05; Figure 3a). In particular, a negative correlation between right BA10 activity during explicit processing of angry faces and Adult BG scores was present in C-carriers, while a positive correlation was present in TT individuals (Figure 3b). This interaction explains the lack of a major genotype effect.

**Figure 3:**
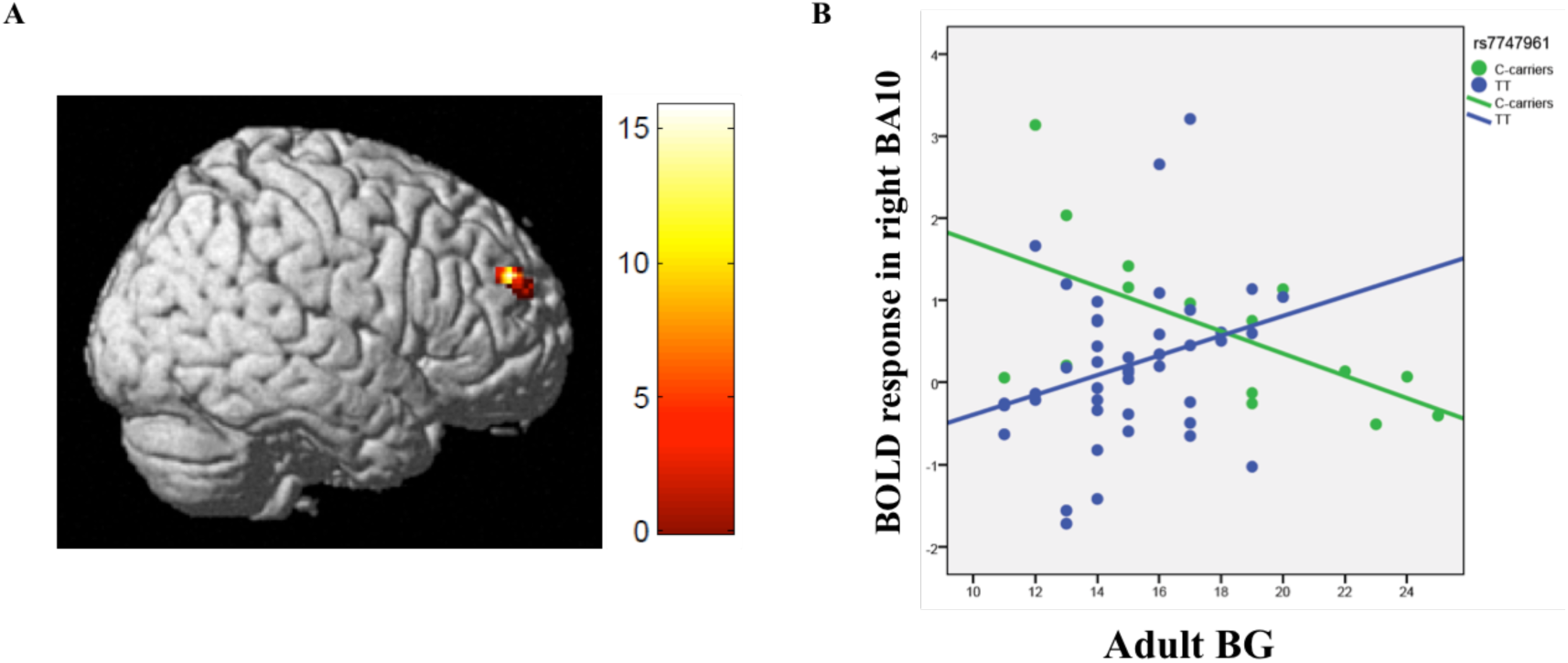
**(A)** Rendered image showing the interaction between Adult BG and rs7747961 in right BA10. **(B)** Scatterplot showing the significantly different relationship in C-carriers and TT subjects between Adult BG and BOLD response in right BA10 during explicit processing of angry faces. See text for statistics.

### Gene co-expression network analysis

Lastly, to gain potential insight into the biological processes related to *LINC01268*, we explored co-expression gene networks that contained *LINC01268*, using the WGCNA algorithm(47). WGCNA was performed on the RNA-seq dataset from the sample of suicide by violent means and identified 10 modules of co-expressed genes. These modules (randomly color-coded for discussion purposes) varied in size from 173 to 1185 genes. Using our conservative approach to RNA quality correction, 16,503 expressed genes were not allocated to any module (“grey” genes). Our module of primary interest (i.e. the module including *LINC01268*) was labeled as “purple” and contained 173 genes. Figure 4 shows the genes within the purple module for which the weights of inter-connectivity (edges) were greater than 0.1, a generally applied threshold. Within the purple module, *LINC01268* had the strongest connection with *P2RY13*, a purinergic receptor present in both the peripheral immune system and in brain(48), while the most connected gene in the whole module, (a.k.a. ‘hub’) was *LAPTM5*, a gene mainly expressed in hematopoietic cells and localized to the lysosome, and shown to modulate inflammatory response by macrophages(49). GO analysis of each module showed statistically significant enrichment in a number of biological processes (BP) (p<0.01, q<0.05 after Benjamini-Hochberg correction) for 8 of the 10 modules. The purple module, containing *LINC01268*, was significantly enriched for 497 GO biological processes gene sets related to immunological functions such as positive regulation of immune response (ST2). Module preservation analysis used to support network validity showed that the purple module was strongly preserved in non-suicide and in suicide by non-violent means expression datasets (SF5, Z_summary_>30 and median rank?1), indicating that *LINC01268* belongs to a network associated with immunity, not limited to suicide by violent means. However, we cannot exclude potential differences between groups in genes connectivity strengths (e.g., differences between groups concerning intra-modular, extra-modular or total connectivity for *LINC01268*), which may be investigated in the future with more detailed analyses(50).

**Figure 4:**
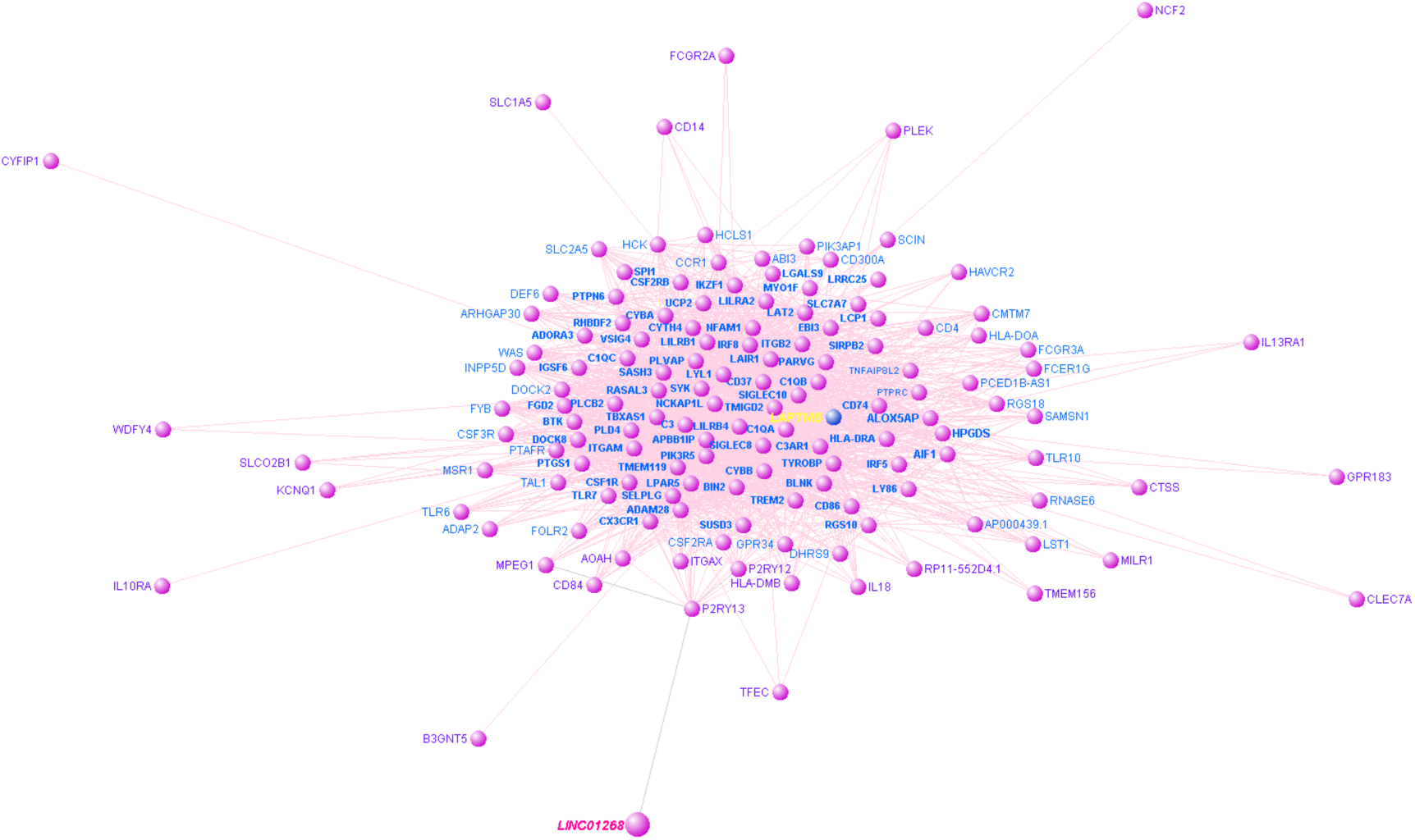
“Purple” module sub-network containing *LINC01268* (figure created with VisANT http://visant.bu.edu/; NB: due to graphical constraints, only genes with strength of connectivity >0.12 are shown, to avoid cluttering). The “hub” (i.e. most connected) gene in the module, *LAPTM5*, is labeled in yellow, and our gene of interest, *LINC01268*, in pink. Within this group of top most connected genes, *LINC01268* has the strongest connectivity only with *P2RY13*.

## DISCUSSION

We report a molecular association in human brain with suicide by violent means that is both strong and replicable, identifying the first non-coding RNA consistently associated with suicidal behavior/aggression. Data from a new sample confirm the presence of a significant difference in DLPFC expression levels of *LINC01268* between suicide and non-suicide deceased, and specifically increased *LINC01268* expression when violent means of suicide are chosen. Moreover, a SNP associated with its expression, rs7747961, is associated with personal ratings of aggressiveness in a cohort of living healthy subjects; the same SNP interacts with a prefrontal physiological assay of processing potentially aggressive stimuli on fMRI.

Finally, *LINC01268* is significantly co-expressed in brain with genes related to the immune response, at least in the periphery.

In contrast to current clinical studies, our findings are from completed suicide cases. Most suicide research looks at survivors, which means studying subjects who did not actually carry out the very behavior being studied (suicide), i.e. who just attempted - or, thought about - various degree of self-harm, in a range of time that may precede the study from weeks to few years. Yet, in an actual suicide, death is the result of willful actions, where the subject hurts him/herself physically and effectively, in the context of a dramatic mental state. In our opinion, that peculiar state of mind, i.e. how that person was feeling at that very moment, may relate to a more salient biology that we can then observe in the brain. We expect that such biology would better stand out from the background when more violent and lethal methods were employed. Indeed, by adopting such precise classification, we were able to strongly replicate the association of a *LINC01268* with suicide by violent means.

The significant increase of *LINC01268* DLPFC expression in suicide compared to non-suicide potentially may offer a tool to differentiate suicidal candidates from patients who, even though affected by the same conditions, will not shift from contemplating death to actually pursuing it. In this regard, levels of *LINC01268* measured in peripheral blood – if linked with brain levels – might represent a further step for *in vivo* studies, leading to prevention and treatment strategies. MARCKS levels in peripheral blood have been reported to show association with suicidal ideation(9). The presence of higher lincRNA expression specifically in suicides by violent means suggests an association of the non-coding RNA with aggressive behaviors, which our data from a questionnaire of aggressive behavior support. The association between an eQTL for the gene with a history of aggressive behaviors in healthy subjects substantiates further the connection and set the stage for our fMRI study.

In the fMRI protocol, the explicit emotional processing of facial expressions of anger was chosen as a proxy for an aggressive stimulus. Interestingly, C-carriers, who are associated with increased *LINC01268* expression in brain and increased aggressiveness, show a negative correlation between DLPFC, specifically Brodmann Area 10 (BA10) activity and aggressiveness, while the opposite is true for TT subjects. BA10 has been implicated in complex cognitive operations(51), and involved in both violent offending(52) and suicide(53–56). A previous report(57), similarly testing anger as a stimulus, also found a negative relationship between aggressive behavior and response of the orbitofrontal cortex (including BA10) in healthy controls, arguing that greater activity facilitates more controlled behaviors. In the present study, we found that this negative relationship between aggressiveness and angry-related BA10 activity (i.e., lower activity with more aggressiveness) is found only in the context of C-carrier rs7747961 genotypes, suggesting that BA10 activity of individuals with major allele TT genotypes are indeed associated with relatively more controlled behavior.

Finally, the biological function of *LINC01268* in brain is largely unknown. To shed light on a potential role, we performed co-expression analysis in brain based on the assumption that genes co-expressed with *LINC01268* represent a molecular network that shares a common function. Weighted network analysis (WGCNA) revealed a *LINC01268-* related network containing genes involved in immunological responses, though such conclusion is based largely on the function of the co-expressed genes in peripheral immune cells, not in brain. Recent evidence points to epigenetic mechanisms and immune and neural pathways as a shared biological background for suicide(58), but whether the immunological mechanisms involved are largely peripheral or central is unclear. Interestingly, the gene most highly correlated with *LINC01268* DLPFC expression is *P2RY13*, which is a recently identified and largely unknown purinergic receptor present in both the peripheral immune system and in brain(48).

In conclusion, our findings converge on implicating a role for *LINC01268* in emotional regulation, aggressive behaviors and suicides by violent means. The biological mechanisms may involve the modulation of genes known to be engaged in immunity and lithium response, and may be relevant from a preventive, diagnostic and therapeutic perspective. Such potential involvement requires testing via *in vitro* or mouse modeling, e.g. by knockdown of the gene coupled with evaluation of molecular regulation of transcription and associated epigenetic signals within the gene locus and distant to it. Future approaches to aggressive self or hetero-directed behaviors might benefit from the targeting of novel and more specific candidates such as those highlighted herein.

## Acknowledgements

We would like to thank Rahul Bharadwaj, Ph.D., for insightful discussions. We are grateful to the Lieber and Maltz families; we are also indebted to all the brave and generous families that consented for the brain donation of their deceased next of kin.

## Notes

The Authors report no financial relationships with commercial interest.

